# Organoid transplantation in the adult endometrium restores fertility and uncovers epithelial lineage plasticity

**DOI:** 10.64898/2026.08.28.747350

**Authors:** Dhanvika Mopure, Hong Im Kim, Claire J. Ang, Daniel J. Davis, Thomas E. Spencer, Kara L. McKinley, Andrew M. Kelleher

## Abstract

The adult endometrium regenerates repeatedly, yet the cells and mechanisms that rebuild its epithelium remain poorly defined. To control the cell types available for regeneration, a genetic model to extensively ablate the uterine epithelium was combined with transplantation of lineage-labeled organoids. Ablation without organoid transplantation triggered re-epithelialization, but resulted in infertility. Transplanted endometrial epithelial organoids engrafted into the ablated uterus, reconstructed both the luminal and glandular epithelia, and restored fertility. Depleting organoids of the glandular lineage before transplantation revealed that luminal epithelial-derived cells acquire glandular identity and function after engraftment. The same luminal-to-glandular epithelial differentiation trajectory emerged during endogenous repair following targeted glandular ablation. Together, these findings establish luminal-to-glandular epithelial conversion as an intrinsic regenerative property of the adult uterine epithelium and establish an endometrial organoid transplantation platform with therapeutic potential.

## Introduction

The adult endometrium before menopause has the capacity for scar-free regeneration, cycling repeatedly through proliferation, differentiation, breakdown, and regeneration to restore full function after menstruation and parturition^1–4^. This intrinsic capacity for regeneration implies an active progenitor population, yet the identity of those cells and the mechanisms governing epithelial regeneration remain poorly defined^3–5^. The endometrial epithelium consists of two major morphologically and functionally distinct cell types, the luminal epithelium (LE) and the glandular epithelium (GE). In both mouse and human development, the GE differentiates from precursor LE cells that bud from the luminal surface and elongate into the underlying stroma^6,7^. Uterine gland development is essential for uterine function, embryo implantation, and placentation^7^. In the adult, GE and LE cells make varying contributions to regeneration depending on species and tissue context, and several epithelial cell types associated with specific marker genes have been proposed as epithelial progenitors^8–12^. During human menstruation, large portions of the LE, GE, and the surrounding stroma are lost, then epithelial cells migrate from remaining LE and residual GE harboring putative endometrial epithelial stem cells to rebuild the luminal surface^3,13–16^. The progenitors responsible for repair after human parturition remain poorly defined, and postpartum repair takes considerably longer than menstruation, reaching completion by approximately six weeks postpartum^17,18^. Interestingly, in mouse models of menstruation and parturition, persistent LE cells may make significant contributions to re-epithelializing the endometrial surface^13^. Directly testing the regenerative potential of defined epithelial populations remains a critical goal for the field^10^.

Endometrial organoid technology provides a platform for studying epithelial biology and for testing these questions directly^19–22^. Epithelial organoid transplantation extends this control into the native tissue and has reconstructed functional epithelium across several organs. This paradigm has been developed most fully in the intestine, where *in vitro* expanded colonic organoids have been shown to integrate into a denuded mucosal surface and form histologically and functionally normal crpyts^23–25^. These studies demonstrated that culture expanded adult epithelium retains full regenerative competence *in vivo*, and coupling transplantation with lineage marking and genome editing makes it possible to read out stem cell behavior directly^25^. In the endometrium, organoids can engraft and contribute to repair in models of damage, yet the lineage dynamics of the regenerated epithelium and the extent of functional rescue remain underexplored, highlighting the need for a tractable, lineage-resolved transplantation model^26,27^.

Here, we established a genetic model of uterine epithelial ablation together with lineage-marked organoid transplantation. Transplanted endometrial epithelial organoids (EEO) engrafted into the uterus, reconstructed both the LE and GE compartments, and restored fertility. Using lineage-marked donor EEO depleted of GE cells before transplantation, we found that LE lineage cells acquired glandular identity after engraftment. This fate transition did not occur in culture, suggesting the adult uterine microenvironment drives LE-to-GE conversion. Targeted gland ablation revealed the same trajectory during endogenous repair. Together, these findings establish LE-to-GE conversion as an intrinsic regenerative property of the adult uterine epithelium and show that organoid transplantation leverages this plasticity to reconstruct the uterus and restore fertility.

## Results

### Endometrial organoid transplantation restores fertility following epithelial ablation

To establish a platform for epithelial reconstitution in the adult endometrium, we generated an epithelium specific ablation model for the uterus by crossing epithelial-specific *Ltf-iCre* mice^28^ with the *Rosa26-iDTR* line^29^. Administration of diphtheria toxin (DT) targeted uterine epithelial cells, resulting in apoptosis of both LE and GE cells (Figure 1A). Cleaved caspase-3 (CC3), a marker of cell death by apoptosis, was localized only to epithelia at 24 hours post-DT treatment (Figure 1A). Histological analysis of the DT-treated uteri revealed almost total loss of both the LE and GE, whereas the underlying stroma was preserved (Figure 1A). Quantification by flow cytometry and histology determined that DT administration reduced the epithelial population of the uterus by approximately 94% compared with vehicle-treated controls (Figure 1A-1B; Figure S1A). Although the luminal surface was re-epithelialized over the denuded stroma by 10 days post-DT, DT-treated mice failed to establish pregnancies during a three-month breeding trial, indicating that resurfacing of the lumen from retained cells was insufficient to rebuild a functional epithelium (Figure 1C and Figure S1B). Having established that uterine epithelial ablation resulted in infertility, we next asked whether transplantation of EEO could restore uterine function. To enable a genetically tractable model, EEO were generated from *Cxcl15^iCre/+^;Rosa26^nTnG^* donor mice^30^ allowing differential labeling of the LE and GE derived lineages. Cultured EEO contained both GFP- (GE lineage) and RFP-labeled (non-GE lineage) epithelial cells, enabling tracing of donor-derived epithelial lineages after transplantation (Figure 1E). The labeled EEO were transferred into the lumen of *Ltf^iCre/+^;Rosa26^iDTR^*recipient uteri one day (24 hours) after DT administration (Figure 1D). At the time of transplantation, RFP-labeled cells comprised approximately 80% and GFP-labeled cells 20% of the EEO population, roughly reflecting the proportions of the LE and GE *in vivo*^7,31–33^ (Figure 1F and Figure S1C).

**Figure 1.**
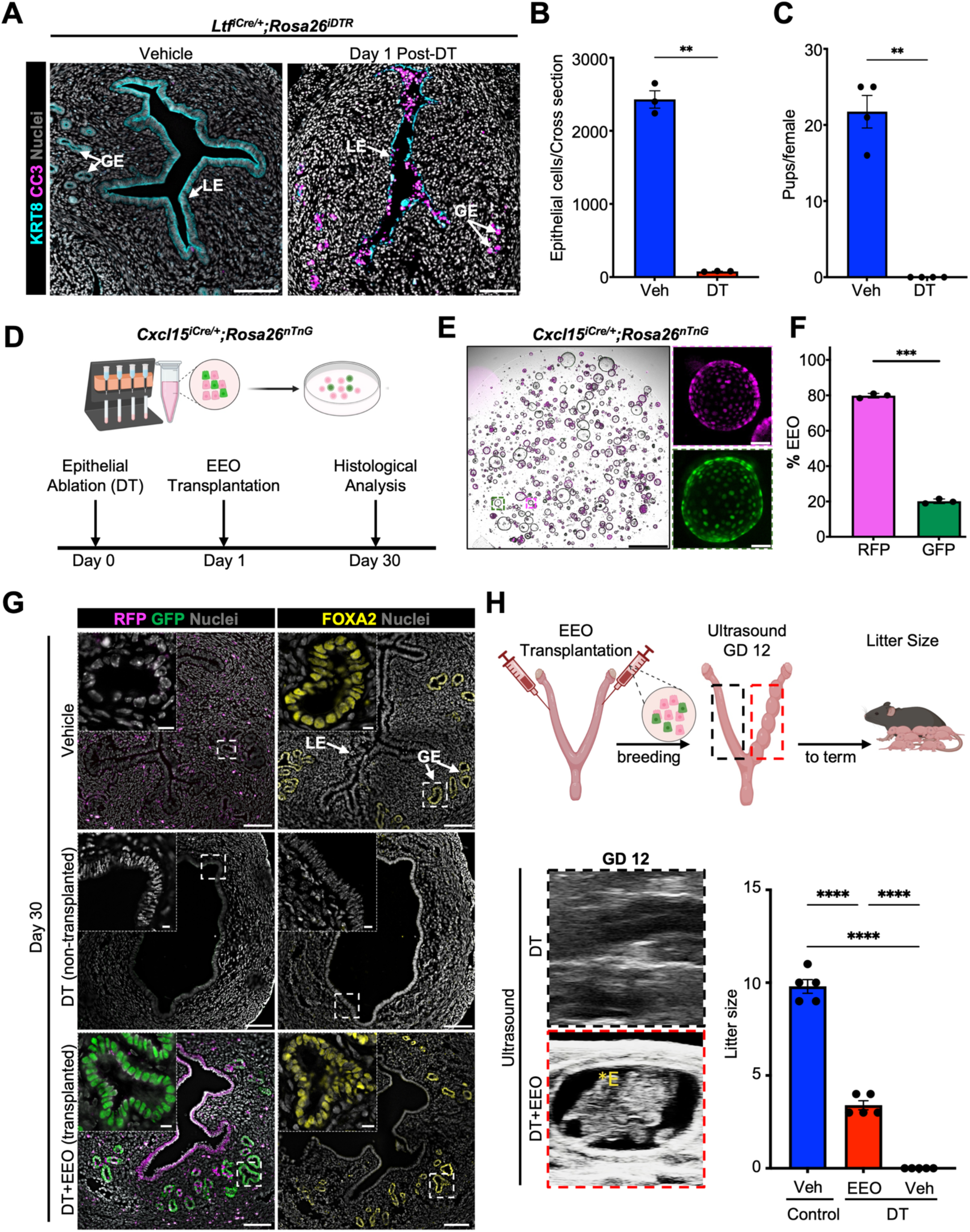
Endometrial organoid transplantation restores fertility following epithelial ablation. (A) Representative immunofluorescence staining for KRT8, Cleaved caspase-3 (CC3) in vehicle and diphtheria toxin (DT) treated *Ltf*^iCre/+^;*Rosa26*^iDTR/+^ on day 1 following DT administration. Scale bar: 100 µm. (B) Quantification of epithelial ablation efficiency in vehicle and DT-treated mice on day 1 following DT administration. Epithelial cells were quantified using Phenoimager analysis of KRT8 immunofluorescence-stained uterine cross sections. *n* = 3 mice per group, with three uteri cross sections analyzed per animal. Statistical analysis was performed using a two-tailed unpaired *t*-test (\*\**P*=0.0025). (C) Total number of pups produced during a three-month fertility assessment of vehicle and DT- treated *Ltf*^iCre/+^;*Rosa26*^iDTR/+^females. *n* = 4 mice per group. Statistical analysis was performed using a two-tailed unpaired t-test (\*\**P*=0.0020). (D) Experimental Schematic illustrating the establishment of endometrial epithelial organoids (EEO) from *Cxcl15*^iCre/+^;*Rosa26*^nTnG^ donor mice, transplantation into DT-treated *Ltf*^iCre/+^;*Rosa26*^iDTR/+^ females on day 1 after epithelial ablation, and uteri collection on day 30. (E) Epifluorescence images showing RFP-and GFP-labeled donor-derived EEOs. Scale bar: 1mm, inset: 50 µm. (F) Flow cytometric quantification of percentage of RFP- and GFP- labeled EEOs at passage 3 from donor mice. *n* = 3 per group. Statistical analysis was performed using a two-tailed paired student’s t-test (\*\*\**P*=0.0006) (G) Epifluorescence images showing RFP and GFP fluorescence together with immunofluorescence staining for GE marker FOXA2 in vehicle and DT-treated *Ltf*^iCre/+^;*Rosa26*^iDTR/+^ on day 30 following transplantation of donor EEOs. Scale bar: 100 µm, inset: 10 µm. (H) Representative B-mode ultrasound image showing an intrauterine embryo in the EEOs-transplanted horn on gestational day (GD)12. Image acquisition was performed using a UHF46X linear probe operating at 47MHz and quantification of litter size in vehicle-treated controls and DT-treated EEOs transplanted and non-transplanted *Ltf*^iCre/+^;*Rosa26*^iDTR/+^ females. *n* = 5 per group. Statistical analysis was performed using a one-way ANOVA (\*\*\*\**P*<0.0001) All uterine cross sections are presented mesometrial (M) to anti-mesometrial (AM) axis. LE, Luminal epithelium; GE, glandular epithelium; Veh, Vehicle; DT, Diphtheria Toxin; EEO, Endometrial Epithelial Organoids. In all graphs, data are presented as mean ± SE, and each dot represents an independent biological replicate. Nuclei were counterstained with Hoechst. Insets show higher magnification.

Thirty days after transplantation, engraftment of donor-derived epithelial cells was observed in transplanted uterine horns. Donor-derived epithelial cells generated LE and GE (Figure 1G and Figure S1D). The GFP-positive cells of the GE expressed FOXA2, and the RFP-positive cells in the LE expressed CALB1 (Figure 1G and Figure S1E), consistent with acquisition of GE and LE identity^7,31,34–36^. These data indicate that transplanted EEO contributed to both the LE and GE compartments of the uterus.

Next, to determine whether EEO transplantation restored uterine function, female recipient mice were mated with fertile males 30 days after transplantation. On gestational day (GD) 12, implantation sites were observed by ultrasound in transplanted uterine horns but not in non-transplanted horns (Figure 1H and Supplemental Video 1). Pregnancies established within transplanted horns progressed to term and resulted in live pups, indicating successful functional engraftment (Figure 1H). The reduction in pups per litter was consistent with pregnancy establishment in only the EEO transplanted horn. Notably, even after pregnancy and postpartum regeneration, donor-derived epithelial cells were retained within the epithelium (Figure S1F). Together, these findings establish that EEO functionally engraft into the adult uterus, contribute to the reconstruction of the LE and GE, and restore uterine function sufficient to support pregnancy.

### Luminal epithelial-derived organoids generate glandular epithelium

Having established functional engraftment with a mixed EEO population, we next leveraged this model to assess the regenerative potential of the adult LE cells and their ability to generate new GE *in vivo*. To test this directly, we generated EEO from triple-transgenic *Cxcl15^iCre/+^;Rosa26^iDTR/nTnG^*mice, in which *Cxcl15*-expressing GE cells express the DTR and are permanently marked by nuclear GFP. To selectively eliminate GFP-positive cells from donor EEO, we treated the *in vitro* EEO cultures with DT, which ablated GFP-labeled GE cells while preserving the RFP-positive LE population (Figure 2A-B). The RFP-positive EEO were transferred into the lumen of epithelial-deficient *Ltf^iCre/+^;Rosa26^iDTR^*recipient uteri 24 hours after DT treatment (Figure 2A). Ablation of the GFP cells in the donor EEO population was confirmed by epifluorescence and flow cytometry, and GFP-positive cells did not re-emerge during extended *in vitro* culture, indicating the purity of the RFP donor population (Figure 2B and Figure S2A).

**Figure 2.**
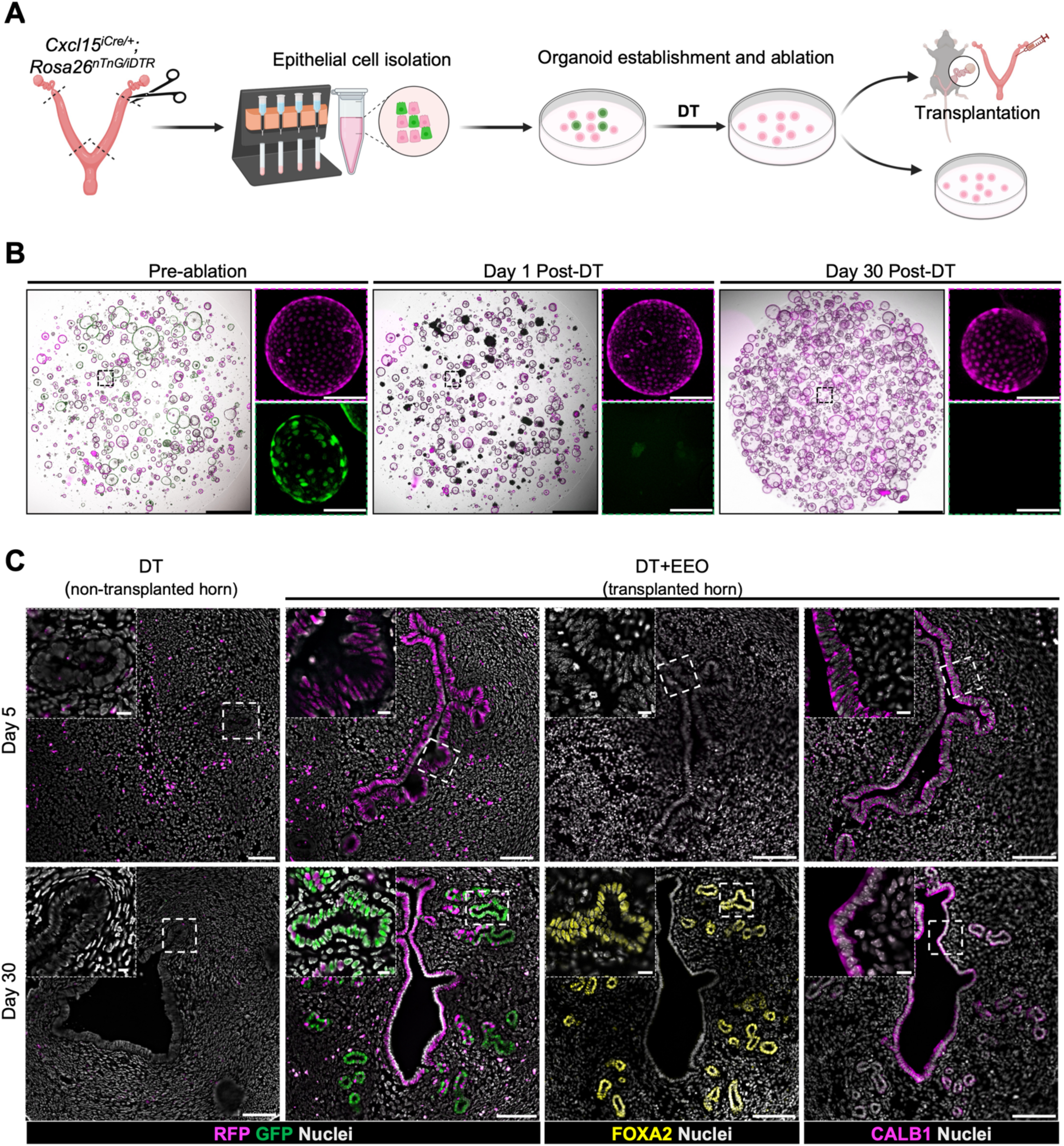
Luminal epithelial-derived organoids generate glandular epithelium following transplantation. (A) Schematic depicting the experimental workflow for establishment and selective depletion of GFP-labeled GE cells in EEOs from *Cxcl15*^iCre/+^;*Rosa26*^iDTR/nTnG^ donor mice and transplantation into the uterine lumen of recipient mice. (B) Brightfield and epifluorescence images of donor-derived EEOs pre-ablation, day 1 post-ablation and day 30 post-ablation in culture. Inset show higher magnification views of single organoid. Scale bar: 1mm, inset: 10 µm. (C) Epifluorescence images showing RFP and GFP fluorescence together with immunofluorescence staining for FOXA2, luminal epithelium (LE) marker CALB1 in vehicle and DT-treated *Ltf*^iCre/+^;*Rosa26*^iDTR/+^ uteri at day 5 and day 30 following organoid transplantation. Scale bar: 100 µm, inset: 10 µm. DT, Diphtheria Toxin; EEO, Endometrial Epithelial Organoids. Nuclei were counterstained with Hoechst. Insets show higher magnification.

At 5 days post-transplantation, histological and immunofluorescence analyses showed that donor-derived epithelial cells remained exclusively RFP-positive, consistent with transplantation of an RFP-positive population (Figure 2C and Figure S2B). By 30 days after transplantation, donor-derived epithelial cells had reconstituted both the LE and GE compartments of the regenerated host endometrium. Cells lining the luminal surface retained RFP expression and were positive for the LE marker CALB1, whereas donor epithelial cells within the glands were GFP-positive, indicating the onset of *Cxcl15-Cre* expression and resulting recombination (Figure 2C). The GFP-positive glands also expressed FOXA2, confirming acquisition of GE identity. This conversion was accompanied by restoration of uterine function to support pregnancy in the transplanted horn(Figure S2C). These findings indicate that EEO lacking GE identity at the time of transplantation acquired glandular fate and functional competence following engraftment into the uterus.

### Functional regeneration of uterine glands following targeted ablation

The transplantation studies found that LE-derived EEO could generate GE, but whether this conversion was an artifact of EEO culture and transplantation remained unknown. To test this, we used the *Cxcl15^iCre/+^;Rosa26^iDTR/nTnG^*mouse model to perform *in vivo* GE ablation. As expected, Cre-mediated recombination resulted in nuclear GFP expression specifically within GE, confirming efficient and restricted recombination. Further, GFP signal was not detected in the LE, stroma, or myometrium, demonstrating lineage specificity (Figure 3A and Figure S3A). Intrauterine administration of DT to *Cxcl15^iCre/+^;Rosa26^iDTR/nTnG^*mice resulted in specific and efficient ablation of GFP-positive GE cells with the apoptosis marker CC3 restricted to the GE and a loss of GFP and FOXA2 protein at 24 hours post-DT (Figure 3B and Figure S3B).

**Figure 3.**
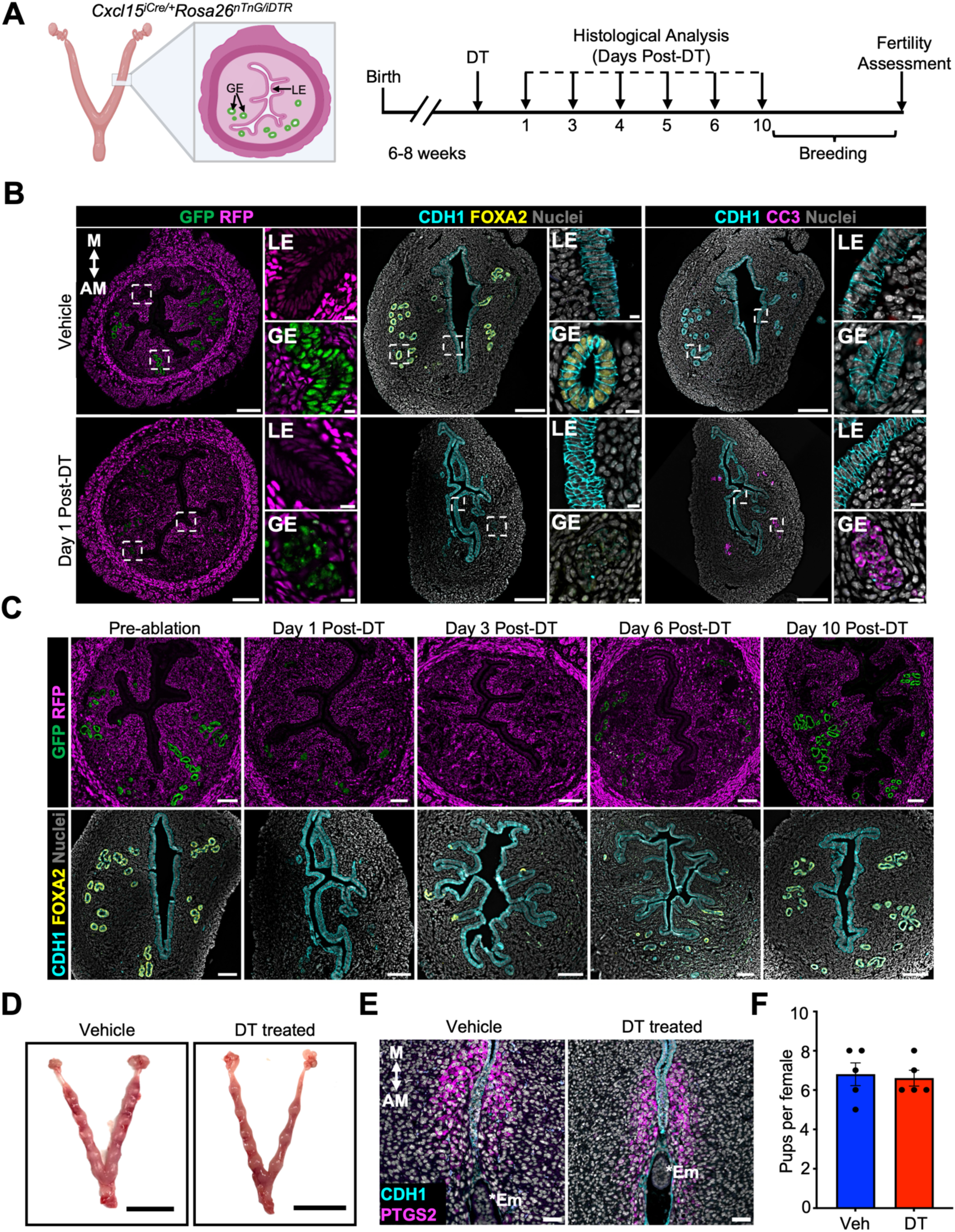
Functional regeneration of the glandular epithelium following ablation. (A) Schematic illustration of a transverse section of adult *Cxcl15*^iCre/+^;*Rosa26*^iDTR/nTnG^ mouse uterus showing that GFP expression is specifically localized to the GE, and an experimental timeline indicating GE ablation, regeneration and functional assessment of regenerated GE. (B) Epifluorescence images showing RFP and GFP fluorescence together with immunofluorescence staining for FOXA2, and Cleaved caspase 3 (CC3) in vehicle and DT-treated *Cxcl15*^iCre/+^;*Rosa26*^iDTR/nTnG^ uteri at day 1 following administration of DT. Scale bar: 100 µm, inset: 10 µm. (C) Epifluorescence images and immunofluorescence staining for FOXA2 during glandular epithelial regeneration at the indicated time points following DT administration. Scale bar: 100 µm. (D) Representative gross images of GD 6 uteri showing implantation sites from vehicle and DT treated female. Scale bar:1 cm. (E) Representative immunofluorescence staining of PTGS2 expression in the GD 6 implantation sites in vehicle and DT treated *Cxcl15*^iCre/+^;*Rosa26*^iDTR/nTnG^ females. Scale bar: 50 µm. (F) Quantification of litter size following fertility assessment is shown *n* = 5 mice per group. Statistical analysis was performed using two tailed unpaired t-test (ns; *P*=0.7854). M, Mesometrial; AM, anti-mesometrial; DT, Diphtheria Toxin; LE, Luminal epithelium; GE, glandular epithelium; EM, Embryo. In all graphs data presented as mean ± SE and each dot represent independent biological replicate. Nuclei were counterstained with Hoechst in all panels. Insets show higher magnification.

To define the regenerative response following GE loss, uteri were analyzed following ablation (Figure 3A). A progressive increase in GFP-positive glandular structures occurred over time, and by day 10 post-DT the endometrial glandular architecture was re-established and composed of FOXA2-positive and GFP- labeled epithelial cells (Figure 3C). To assess the functional competence of regenerated glands, GE-ablated and control females recovered for 10 days and were mated with fertile males (Figure 3A). Both vehicle-treated and DT-treated mice exhibited normal mating behavior, as indicated by the presence of copulatory vaginal plugs. At GD 6, there were no differences in implantation site number observed in the GE-ablated mice (Figure 3D and Figure S3C). Embryos in DT-treated uteri were correctly positioned on the antimesometrial side of the uterus, with removal of the LE adjacent to the implanting embryo^37,38^. Likewise, stromal cells underwent decidualization in both control and regenerated uteri and expressed PTGS2, a marker of primary stromal decidualization^37,39,40^ (Figure 3E). The DT-treated mice were able to produce live pups, and litter sizes not different from controls (Figure 3F), indicating that GE loss was followed by restoration of glandular architecture and function, with regenerated glands expressing FOXA2 and *Cxcl15* and supporting normal implantation, decidualization, and fertility.

### Luminal epithelium regenerates glands through a stepwise transition

To define the cellular origin and differentiation dynamics during gland regeneration, we collected uteri from *Cxcl15^iCre/+^;Rosa26^iDTR/nTnG^*mice at day 3, day 6, and day 10 following DT-mediated GE ablation. At day 3 after ablation, epithelial structures stemming from the LE and lacking GFP expression were observed in the endometrial stroma. Those cells had varieagated FOXA2 expression and a flattened morphology characteristic of invaginating epithelial buds extending from the luminal epithelium into the underlying stroma. The newly formed GE-like structures retained expression of the LE marker CALB1^31,36,41^, demonstrating that early regenerating cells maintained LE characteristics during the initial stages of gland regeneration (Figure 4A). Adjacent LE cells were Ki67 positive, indicating active proliferation during regeneration (Figure S4A). By day 6 after ablation, GFP-positive epithelial cells began to emerge in the developing GE, consistent with a transitional epithelial population acquiring glandular identity in a stepwise manner (Figure 4A). By day 10, the regenerated endometrium contained FOXA2-positive and GFP-positive glandular structures. In contrast to the earlier stages, those regenerated glands were no longer positive for CALB1, demonstrating loss of LE characteristics and acquisition of a mature glandular epithelial phenotype (Figure 4A). These observations indicate a stepwise regenerative process in which the LE cells contribute to glandular regeneration following injury. The progression from CALB1-positive epithelial buds that lack GFP expression, to FOXA2-positive, and finally to mature GFP-postive glands lacking CALB1 supports a model in which LE cells undergo lineage conversion to re-establish the GE (Figure 4B).

**Figure 4.**
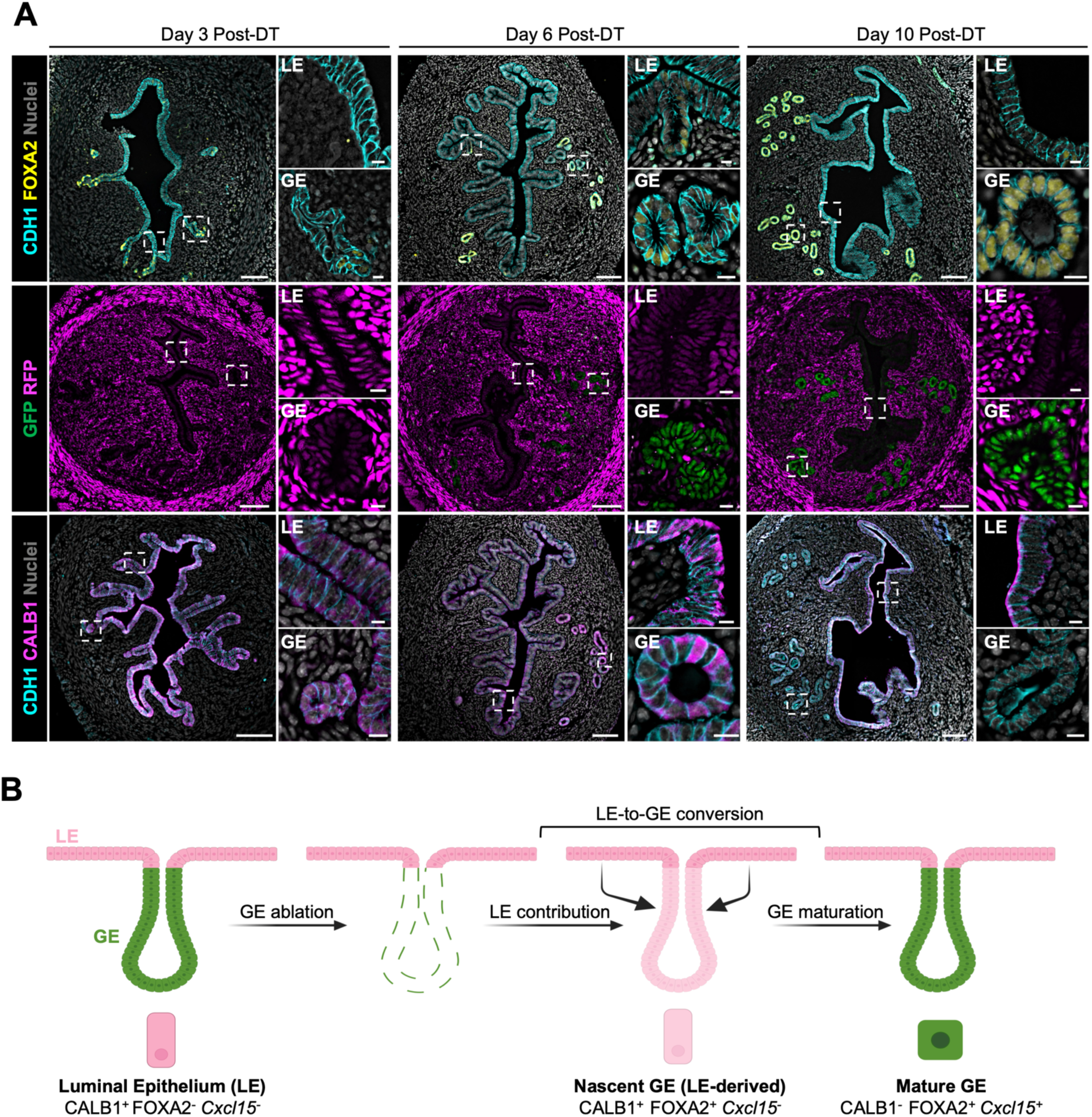
Luminal epithelial differentiation following glandular ablation. (A) Epifluorescence images showing RFP and GFP fluorescence and immunofluorescence staining for FOXA2 and CALB1 during glandular epithelial regeneration at day 3, day 6 and day 10 following DT administration. Nuclei were counterstained with Hoechst. Scale bar: 100 um, inset: 10 um. (B) Schematic of stepwise GE regeneration depicting loss of GE following administration of DT and contribution of *Cxcl15-* negative epithelial cells to glandular epithelial regeneration. DT, Diphtheria Toxin; LE, Luminal epithelium; GE, glandular epithelium.

## Discussion

Despite the importance of uterine regeneration for reproductive health, disease, and fertility, the cellular mechanisms that underlie it remain mostly unresolved. Foundational histological work, mouse genetic models, and more recent transcriptomic studies have proposed several candidate epithelial progenitor populations in the endometrium, most prominently cells residing in the endometrial glands^4,11,42–44^, as well as contributions from bone marrow-derived cells^45^, mesenchymal-to-epithelial transition^46^, and perivascular cells^47^. However, no single model of endometrial re-epithelialization has been widely adopted, and these proposed sources are not necessarily mutually exclusive. Progress has been limited in part by a lack of tractable systems in which the functional regenerative potential of defined epithelial populations can be directly tested.

Organoid models have provided a means to interrogate the regenerative potential and stem-like function of the endometrial epithelium and to identify candidate regulators of these processes^22^. Here, we extended these models by performing *in vivo* transplantation to accelerate our understanding of endometrial lineage dynamics. Coupling targeted epithelial ablation with organoid engraftment and a dual-reporter donor strategy enabled functional assessment of donor organoids *in vivo* anddemonstrated their capacity to contribute to both the LE and GE lineages and restore fertility. A central finding of our study is that the LE cells can generate GE in the adult uterus. In both the engineered transplantation setting and following *in situ* gland ablation, LE-derived cells gave rise to *Cxcl15*-expressing, FOXA2-positive glands. Reciprocally, Ang and coworkers demonstrated the ability of *Cxcl15-*expressing GE to functionally repopulate the epithelial lining of the uterus following chemical ablation^13^. These independent systems indicate that interconversion of epithelial lineages is an inherent inducible property of the adult uterine epithelium, likely dependent on instructive cues from the underlying stroma ^48,49^

The transplantation assay used here allowed for the study of epithelial lineage dynamics in the regenerating uterus, and together with *in vitro* cell-type ablation, our data indicate that LE-derived organoids can reconstitute the ablated epithelial lining. Bipotent cells have been proposed at the intersection of the GE and LE in the mouse^50^, and more recently in the human endometrium between the basalis and functionalis^44^; such cells may be among the *Cxcl15*-negative population that gives rise to new glands. However, this model does not readily account for the capacity of the *Cxcl15*-positive GE population to fully reconstitute the LE^13^. An underexplored alternative is that the endometrial epithelium harbors facultative progenitors, analogous to those described in the liver, intestine, and other epithelia, in which differentiated cells retain the latent competence to re-enter a progenitor-like state upon injury^51–53^.

Our *in situ* glandular ablation model demonstrates the dynamics of this conversion during endogenous repair, in which a stepwise trajectory of CALB1-positive LE cells gives rise to FOXA2-positive epithelial buds that then develop through a transitional state to mature into CALB1-negative, *Cxcl15*-positive GE. Because the GE arises from the LE during early postnatal uterine development^7,31^, this progression suggests that a latent developmental program is reactivated following injury in the adult uterus, as has been described in other regenerating organs^51,54^.

In summary, we establish a transplantation strategy that enables engraftment and a functional readout of the contribution of defined endometrial cell types to regeneration. By demonstrating the functional differentiation potential of the LE and GE, our results provide a foundation for the mechanistic dissection of the pathways that underlie endometrial regeneration during homeostasis and in disease. Looking forward, organoid transplantation not only enables fundamental discovery but also has translational potential^23–25^. For example, niche-preserving engraftment of intestinal organoids functionally restores the epithelium in a rat model of short bowel syndrome^24^. Further evaluation of endometrial epithelial reconstitution as a therapeutic strategy for restoring fertility in cases of epithelial dysfunction is an important direction for future work.

### Limitations

The manuscript is epithelial-focused and does not decipher the stromal, immune, and vascular contributions that accompany endometrial regeneration. The specific niche signals that underlie LE to GE fate transition remain undefined. Our lineage analyses rely on *Cxcl15*-based labeling of the GE. Although recombination was restricted to the glandular compartment, we cannot formally exclude rare labeling of non-glandular cells. Finally, these findings are established in the mouse, and their relevance to human endometrial regeneration and to organoid-based therapeutic strategies will require further investigation.

## Methods

All key resources in this study are listed Table S1.

### Animals

All experimental procedures were approved by the University of Missouri Institutional Animal Care and Use Committee and were performed in accordance with the NIH Guide for the Care and Use of Laboratory Animals. *Cxcl15*^iCre^ mice were generated in house^30^. *Rosa26*^nTnG^ (nuclear tdTomato-to-nuclear EGFP Cre reporter; stock no. 021309), *Rosa26*^iDTR^ (Cre-inducible diphtheria toxin receptor; stock no. 007900), and *Ltf*^iCre^ (lactoferrin-iCre, epithelial-specific; stock no. 026030) mice were obtained from The Jackson Laboratory. All strains were maintained on a C57BL/6J background and housed under a 12 h:12 h light:dark cycle with food and water available ad libitum. Adult females aged 8 to12 weeks of age were used for all experiments unless otherwise noted.

The following genotypes were used. For endometrial epithelial ablation, *Ltf*^iCre/+^;*Rosa26*^iDTR/+^ females were compared with Cre-negative wild-type (WT) littermate controls. For selective GE ablation and for lineage-marked organoid donors, *Cxcl15*^iCre/+^;*Rosa26*^iDTR/nTnG^ triple-transgenic females were used. Dual-reporter donor EEO without the *iDTR* allele were derived from *Cxcl15*^iCre/+^;*Rosa26*^nTnG^ females.

### Diphtheria toxin-mediated epithelial ablation

For endometrial epithelial ablation, *Ltf*^iCre/+^;*Rosa26*^iDTR/+^ females and WT controls received a single intraperitoneal injection of diphtheria toxin (DT; Cayman Chemical, 19657) at 25 µg/kg body weight in phosphate-buffered saline (PBS). For selective glandular epithelial ablation, *Cxcl15*^iCre/+^;*Rosa26*^iDTR/nTnG^ females and WT controls received a single uterine intraluminal injection of 5ng DT in 15 µL of 10% Cultrex in PBS into one uterine horn. Tissues were collected at indicated time points after DT administration (n = 4-6/timepoint).

### Mouse endometrial epithelial cell isolation and organoid establishment

EEO were established and cultured using established protocols^55–58^. Oviducts, uterine horns, and cervices were dissected from donor females. Uterine horns were cut into three pieces and digested in Liberase (0.2 mg/mL; Roche, 5401119001) in phenol red-free RPMI 1640 supplemented with DNase-I (0.5 mg/mL; Roche, 10104159001) on an orbital shaker for 45 min at 4°C followed by 45 min at 37°C. Digestion was stopped with base organoid medium (Table S2). Epithelial and stromal fractions were separated by two rounds of magnetic sorting with CD326 (EpCAM) MicroBeads according to the manufacturer’s instructions (Miltenyi Biotec, 130-105-958). EpCAM^+^ epithelial cells retained on the magnetic column were eluted, centrifuged (300 × g, 5 min), resuspended in Cultrex (R&D Systems, 3433-001-R1), and (plated in 12-well plates 20 µL drops, 4 drops per well). Cultrex drops were solidified at 37°C for 15 min before addition of 800 µL organoid expansion medium per well (Table S3).

Organoids were passaged every 7–10 days at a 1:3 ratio. Cultrex drops were dislodged by pipetting, collected, centrifuged (300 × g, for 3 minutes), dissociated in base organoid medium, re-pelleted, resuspended in Cultrex, and replated as above. All transplantation and imaging experiments used EEO from passage 3-5. Brightfield and epifluorescence images were captured on a Leica DMi8 inverted microscope with a Leica K8 camera using Leica Application Suite X (LAS X) (n= 4-12 drops/biological replicate).

### Depletion of glandular epithelial cells from donor EEO

Passage two *Cxcl15^iCre/+^;Rosa26^iDTR/nTnG^*EEO were treated with Diphtheria Toxin (DT; 1 ng/mL of media) to ablate GFP-positive cells while sparing the RFP-positive luminal epithelial cell population. Depletion of GFP cells was confirmed by epifluorescence and flow cytometry.

### Fluorescence-activated cell counting

For quantification of EEOs, organoids were collected at passage 3 and dissociated to single cells with TrypLE (Gibco, 12563011) supplemented with DNase-I and ROCK inhibitor (PeproTech, 1293823). Cells were resuspended in a neutral buffered solution (PBS, 2% FBS, 2 mM EDTA) and filtered through a 40–70 µm cell strainer. Single cell suspensions were analyzed on a Cytek-Aurora flow cytometer (University of Missouri Flow Cytometry Core). The gating strategy was as follows: all events (singe cells) > GFP^+^ or RFP^+^ cells.

For quantification of epithelial content in *Ltf*^iCre/+^;*Rosa26^iDTR/+^*on day 1 post-DT administration, isolated cells were collected in a neutral buffered solution (PBS, 2% FBS, 2 mM EDTA), filtered through a 40–70 µm cell strainer and stained with anti-CD326 (EpCAM; PE; Invitrogen, 12-5791-82) for 30 min at 4°C. DAPI was used to identify and exclude non-viable cells. The stained cells were washed and analyzed on the Cytek-Aurora. The gating strategy was as follows: All events (single cells) > DAPI negative viable cells > EpCAM (PE)^+^ cells. Unstained and single-stained references were used for compensation/unmixing controls.

### Uterine epithelial cell transplantation

Recipient *Ltf*^iCre/+^;*Rosa26^iDTR/+^* females received DT as described above and received organoid transplantation 24 hrs later. A suspension of 1 million cells derived from EEO was prepared in 15 µL of Cultrex (20 µL total), and for vehicle controls, 5 µL of DPBS was suspended in 15 µL of Cultrex immediately before surgery. Recipients were anesthetized with 3% isoflurane for induction and maintained at 2.5% isoflurane. The flanks were shaved and aseptically prepared with alternating betadine and 70% ethanol. A small dorsal skin incision was made above the hind limb, followed by an incision through the body wall. The ovarian fat pad was exteriorized to expose the ovary and uterine horn. Prior to injection, mini occlusion clips were applied to the ovarian and cervical ends of uterine horn to prevent reflux of the injected cells. A 30-gauge needle was inserted into the proximal uterine horn immediately distal to the utero-tubal junction, and the cell suspension was slowly injected along the length of the uterine lumen. The horn was held for 45 to 60 seconds to facilitate the retention of injected cells, after which the clips were removed and the horn was returned to the abdominal cavity. The body wall was closed with absorbable suture and the skin with a wound clips. The contralateral uterine horn received vehicle control (5 µLof DPBS in 15 µLof Cultrex).

### Fertility assessment

Control and experimental females were housed individually with proven-fertile CD-1 males, and litter sizes were recorded over the duration of the trial. For timed collections, mating was confirmed by the presence of a copulatory plug, designated gestational day 1 (GD1). On GD6, implantation and inter-implantation sites were dissected separately; pregnancy was additionally assessed by ultrasound on GD12 (n = 3-5 mice per treatment)

### Ultrasound imaging

Pregnancy was assessed non-invasively at GD12 on a Vevo F2 LAZR-X small-animal ultrasound system (FUJIFILM VisualSonics). Mice were anesthetized with 3% isoflurane and positioned supine on a heated stage. A 47 MHz transducer (MS550D-0421) was mounted on a motorized stage, B-mode image stacks were acquired across the abdomen in 10 µm steps and reconstructed and analyzed in three dimensions with Vevo Lab software (version 5.5.1).

### Histological quantification of epithelial cell ablation

Epithelial cells were quantified on immunofluorescence-stained uterine cross-sections from vehicle- and DT-treated animals using PhenoImager analysis software (Akoya Biosciences). Whole-slide images were acquired with identical settings across samples. Regions of interest encompassing the uterine epithelium were defined, nuclei were segmented on DAPI, and epithelial cells were scored as DAPI-positive nuclei within the KRT8-positive epithelial compartment. Identical thresholding and segmentation parameters were applied across all groups. The total epithelial cell count per cross-section was used for statistical comparison (3 cross-sections per animal from three females).

### Histology and immunofluorescence

Uteri were fixed in 4% paraformaldehyde in PBS overnight at 4°C, dehydrated, paraffin-embedded, and sectioned at 5 µm. Sections were baked for 30 min at 60°C, deparaffinized in xylene, and rehydrated through graded ethanol. Antigen retrieval was performed in 10 mM citrate buffer (pH 6.0) at 95°C for 15 min followed by cooling to room temperature and permeabilized with 0.2% Triton X- 100 in PBS for 10 min. Sections were blocked in 5% (v/v) normal goat serum in PBS (pH 7.2) for 30 min and incubated with primary antibodies overnight at 4°C. The following primary antibodies were used: rabbit monoclonal anti-FOXA2 (1:800; Abcam, ab108422), rabbit polyclonal anti-Ki67 (1:500; Abcam, ab15580), rabbit monoclonal anti-COX2 (PTGS2) (1:500; Abcam, ab179800), rat monoclonal anti-cytokeratin 8 (1:200; EMD Millipore, MABT329), rabbit polyclonal anti-cleaved caspase-3 (1:500; Cell Signaling Technology, 9661), mouse anti-E-cadherin (1:500; BD Biosciences, 610182), and rabbit monoclonal anti-calbindin 1 (1:800; Cell Signaling Technology, 13176T). Signal was detected with Alexa Fluor 488-, 594-, or 647-conjugated secondary antibodies (1:500; Jackson ImmunoResearch, 112-545-143, 115-585-46, 111-605-144) for 60 min at room temperature. Nuclei were counterstained with Hoechst 33342 (2 µg/mL; Invitrogen, H3570) and sections were mounted in ProLong Diamond Antifade Mountant (Invitrogen, 36961). Images were acquired on a Leica DM6 B upright microscope with a Leica K8 camera using Leica Application Suite X (LAS X).

For cryosections, slides were dried at 45°C for 30 min, permeabilized with 0.2% Triton X-100 in PBS for 10 min, blocked in 5% normal goat serum (NGS), and incubated with primary antibodies against cytokeratin 8 (TROMA-1; 1:200; Developmental Studies Hybridoma Bank, University of Iowa) and FOXA2 (1:700; Abcam, ab108422) overnight at 4°C. Slides were washed in PBS and incubated with Alexa Fluor-conjugated goat anti-rabbit and anti-rat secondary antibodies for 60 minutes at room temperature. Nuclei were stained and slides cover slipped as described above. Images were acquired on a Leica DM6 B upright microscope using Leica Application Suite X (LAS X).

## Quantification and statistical analysis

All mouse and cell culture experiments were performed using at least three biological and technical replicates. Statistical analysis was performed in GraphPad Prism 9 (v.11.0.2). A two-tailed student’s t-test with Welch correction was used for all comparisons between two conditions. In cases with more than two conditions a one-way ANOVA was used, or as described in the figure legends. A *p*-value of less than 0.05 was considered statistically significant.

## Acknowledgements

We thank members of the Kelleher and McKinley labs for their feedback on the manuscript. This work was supported by the National Institutes of Health (R01HD112315 to AMK, R37HD114609 to TES, DP2HD111708 and R00HD101021 to KLM), New York Stem Cell Foundation (KLM), and Howard Hughes Medical Institute (KLM). KLM is a New York Stem Cell Foundation Robertson Stem Cell Investigator. This material is based on work supported by the National Science Foundation Graduate Research Fellowship Program to CJA under grant number DGE 2140743.

