## supplementary materials for "Organoid transplantation in the adult endometrium restores fertility and uncovers epithelial lineage plasticity"

**This PDF file includes:**

Figures S1 to S4
Tables S1 to S3
Video S1 (description)

#### S. Figure 1.

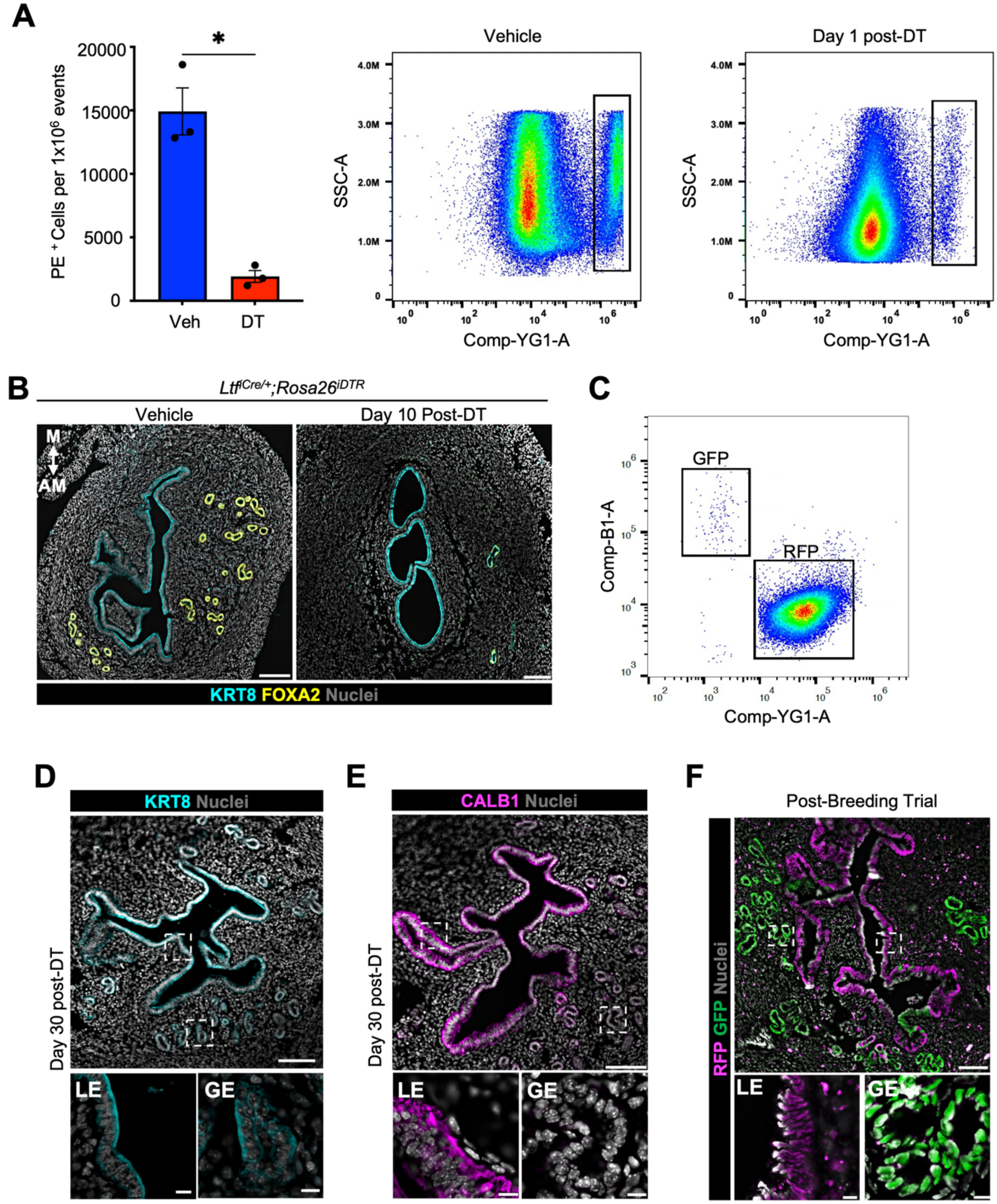
A
PE + Cells per 1x10^6 events
20000
15000
10000
5000
0
*
Veh
DT
Vehicle
Day 1 post-DT
SSC-A
Comp-YG1-A
B
LtfiCre/+;Rosa26iDTR
Vehicle
Day 10 Post-DT
M
AM
KRT8 FOXA2 Nuclei
C
GFP
RFP
Comp-B1-A
Comp-YG1-A
D
KRT8 Nuclei
Day 30 post-DT
LE
GE
E
CALB1 Nuclei
Day 30 post-DT
LE
GE
F
Post-Breeding Trial
RFP GFP Nuclei
LE
GE

**Supplementary Figure 1. Endometrial epithelial ablation and regeneration, related to Figure 1.**

(A) Flow cytometry quantification of PE-positive cells in vehicle and Day 1 diphtheria toxin (DT) injection. *n* = 3 per group. Statistical analysis was performed using two tailed unpaired t-test (* *P*=0.0152) and representative gating strategy to quantify $PE^+$ cells.

(B) Representative immunofluorescence staining for KRT8, FOXA2 in vehicle and DT treated $Ltf^{iCre/+}$;$Rosa26^{iDTR/+}$ on day 10 following DT administration. Scale bar: 100 μm, inset: 10 μm.

(C) Representative flow cytometry plot showing the gating strategy used to quantify percentage of RFP- and GFP- labeled EEOs at passage 3 from donor mice.

(D) Representative immunofluorescence staining for KRT8 on day 30 following EEOs transplantation. Scale bar:100 μm, inset: 10 μm.

(E) Representative immunofluorescence staining for CALB1 on day 30 following EEOs transplantation. Scale bar:100 μm, inset: 10 μm.

(F) Representative epifluorescence images showing RFP and GFP labeled epithelial cells post fertility assessment. Scale bar: 100 μm, inset: 10 μm.

In all graphs, data are presented as mean ± SE, and each dot represents an independent biological replicate. Nuclei were counterstained with Hoechst. Insets show higher magnification.

#### S Figure 2.

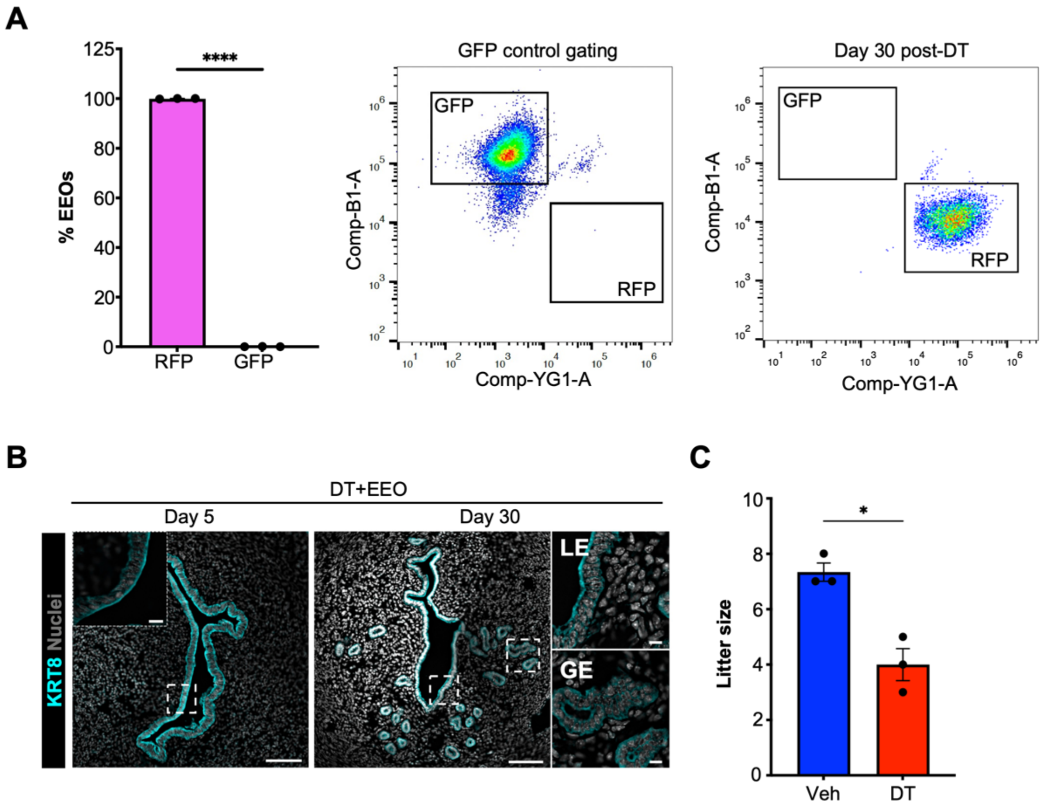


**Supplementary Figure 2. Depletion of GE-derived EEO and functional EEO engraftment, related to Figure 2.**

(A) Flow cytometry quantification of percentage of RFP- and GFP- labeled EEOs from donor mice following selective depletion of GE cells. *n* = 3 per group. Statistical analysis was performed using two tailed paired student's t-test (*****P*<0.0001). And representative gating strategy for identification of RFP- and GFP- labelled cells.

(B) Representative immunofluorescence staining for KRT8 on day 5 and day 30 following EEOs transplantation. Scale bar:100 μm, inset: 10 μm

(C) Total number of pups produced during fertility assessment of vehicle and DT treated following LE EEO transplantation into $Ltf^{iCre/+}$;$Rosa26^{iDTR/+}$females. *n* = 3 mice per group. Statistical analysis was performed using two tailed unpaired t-test (**P*=0.0132).

In all graphs data presented as mean ± SE and each dot represent independent biological replicate. Nuclei were counterstained with Hoechst. Insets show higher magnification.

#### S. Figure 3.

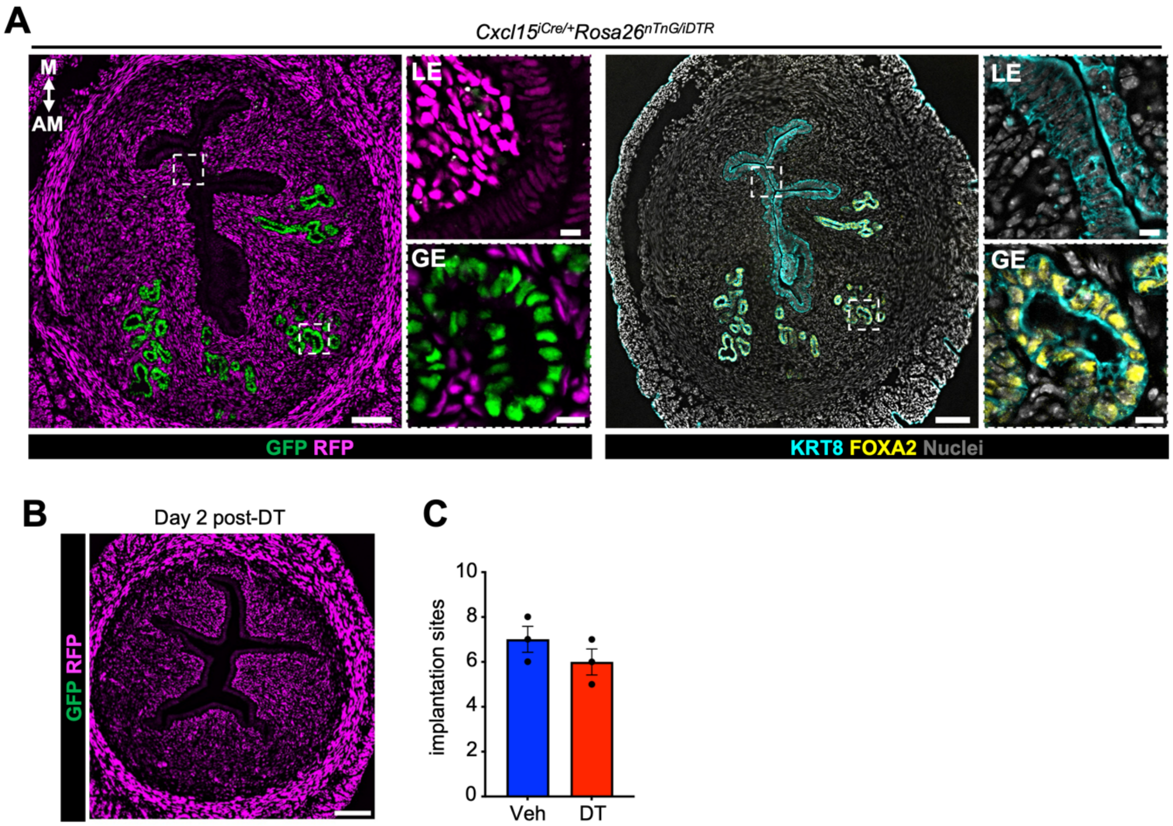


**Supplementary Figure 3. Functional regeneration following *in situ* GE ablation, related to Figure 3.**

(A) Representative epifluorescence image of transverse section of adult *Cxcl15*$^{iCre/+}$;*Rosa26*$^{iDTR/nTnG}$ mouse uterus showing that GFP expression is specific to the GE cells. And immunolocalization of FOXA2 confirms GFP localization to GE. Scale bar: 100 µm, inset: 10 µm.

(B) Representative epifluorescence image showing loss of GFP labelled GE cells by day 2 post-DT. Scale bar: 100 µm.

(C) Quantification of embryo implantation sites on GD 6. *n* = 3 mice per group. Statistical analysis was performed using two tailed unpaired student's *t*-test (ns; *P*=0.2879).

In all graphs data presented as mean ± SE and each dot represent independent biological replicate. Nuclei were counterstained with Hoechst. Insets show higher magnification.

### S. Figure 4.

A

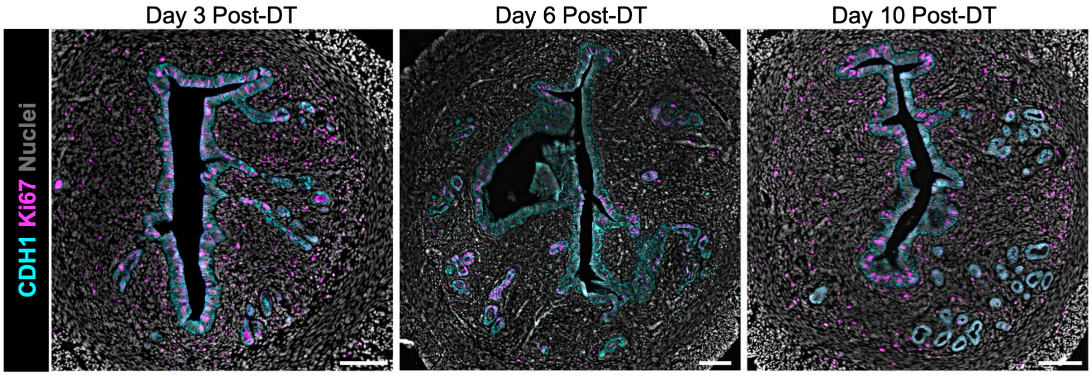


**Supplementary Figure 4. Epithelial proliferation following GE ablation, related to Figure 4.**

(A) Representative immunofluorescence staining for CDH1, Ki67 on day 3, day 6 and day 10 post-DT administration. Nuclei were counterstained with Hoechst. Scale bar:100 µm.

#### Table S1. Key Resources

| REAGENT or RESOURCE | SOURCE | IDENTIFIER |
|---|---|---|
| Antibodies | | |
| Mouse monoclonal anti-CDH1 | BD Biosciences | Cat#610182; RRID: AB_397581 |
| Rabbit monoclonal anti-FOXA2 | Abcam | Cat#ab108422; RRID: AB_11157157 |
| Rabbit polyclonal anti-Ki67 | Abcam | Cat#ab15580; RRID: AB_443209 |
| Rabbit monoclonal anti-COX2 (PTGS2) | Abcam | Cat#ab179800; RRID: AB_2894871 |
| Rabbit polyclonal anti-cleaved CASP 3 (ASP175) | Cell Signaling | Cat#9661; RRID: AB_2341188 |
| Rat monoclonal anti-cytokeratin 8 (TROMA-1) | EMD Millipore | Cat#MABT329; RRID: AB_2891089 |
| Rabbit monoclonal anti-Calbindin 1 | Cell Signaling | Cat#13176; RRID: AB_2687400 |
| Alexa Fluor 594, goat anti-rabbit IgG | Jackson ImmunoResearch | Cat#111-585-144; RRID: AB_2307325 |
| Alexa Fluor 790, goat anti-rabbit IgG | Jackson ImmunoResearch | Cat#111-655-144; RRID: AB_2338086 |
| Alexa Fluor 488, goat anti-mouse IgG | Jackson ImmunoResearch | Cat#115-545-146; RRID: AB_2307324 |
| Alexa Fluor 488, goat anti-rat IgG | Jackson ImmunoResearch | Cat#112-545-143; RRID: AB_2338361 |
| Alexa Fluor 647, goat anti-rabbit IgG | Jackson ImmunoResearch | Cat#111-605-144; RRID: AB_2338078 |
| Anti-mouse CD326 (EpCAM), clone G8.8,PE | Invitrogen | Cat#12-5791-82; RRID: AB_953615 |
| Biological samples | | |
| Murine-derived endometrial epithelial organoids | Mice housed at University of Missouri vivarium | This paper |
| Chemicals, peptides, and recombinant proteins | | |
| Advanced DMEM/F12 | Gibco | Cat#12634028 |
| B27 Supplement | Gibco | Cat#17504044 |
| Insulin-transferrin-selenium | Gibco | Cat#41-400-045 |
| Primocin | InvivoGen | Cat#ant-pm-05 |
| Glutamax | Gibco | Cat#35050061 |
| A83-01 | BioGems | Cat#9094360 |
| Murine EGF | R&D Systems | Cat#2028-EG-200 |
| Murine FGF-10 | PeproTech | Cat#450-61 |
| Murine R-spondin1 | R&D Systems | Cat#7150-RS-010/CF |
| Murine Noggin | R&D Systems | Cat#997-NG-025/CF |
| Murine Wnt3a | R&D Systems | Cat#1324-WN-002/CF |
| Nicotinamide | Millipore Sigma | Cat#N0636-100G |
| N2 | Gibco | Cat#17502001 |
| DPBS | Gibco | Cat#14190144 |
| Liberase | Millipore Sigma | Cat#05401020001 |

| | | |
|---|---|---|
| DNase I | Roche | Cat#10104159001 |
| RPMI | Gibco | Cat#11835030 |
| Cultrex BME | R&D Systems | Cat#3433-010-R1 |
| Y-27632 dihydrochloride | BioGems | Cat#1293823 |
| EPCAM microbeads | Miltenyi Biotec | Cat#130-105-958 |
| Diphtheria toxin (unnicked) | Cayman Chemicals | Cat#19657 |
| Hoechst 33342 | Invitrogen | Cat#H3570 |
| ProLong Diamond antifade mountant | Invitrogen | Cat#36961 |
| Paraformaldehyde (96%) | Thermo Scientific | Cat#AC416780030 |
| Normal goat serum | Invitrogen | Cat#01-6201 |
| Drierite (desiccant) | Fisher | Cat#089751.P5 |
| TrypLE, no phenol red | Gibco | Cat#12563011 |
| Triton X-100 | Thermo Scientific | Cat#AAA16046AP |
| Viastain AOPI staining solution | Revvity | Cat#CS2-0106-25ML |
| Experimental models: Organisms/strains | | |
| *Cxcl15*$^{iCre/iCre}$ | Kelleher et al. 2025 | NA |
| *Rosa26*$^{nT\text{-}nG/nT\text{-}nG}$ | Jackson Laboratory | Cat#021309; RRID: IMSR_JAX:021309 |
| *Rosa26*$^{iDTR/iDTR}$ | Jackson Laboratory | Cat#007900; RRID: IMSR_JAX:007900 |
| C57BL/6J | Jackson Laboratory | Cat#000664; RRID: IMSR_JAX:000664 |
| *Ltf*$^{iCre/iCre}$ | Jackson Laboratory | Cat#026030; RRID: IMSR_JAX:026030 |
| Software and algorithms | | |
| Leica Application Suite X (LAS X) software | Leica | https://www.leica-microsystems.com/products/microscope-software/p/leica-las-x-ls/ |
| GraphPad Prism | GraphPad | https://www.graphpad.com/ |
| Other | | |
| EASYstrainer (70 µm) | Greiner Bio-One | Cat#542070 |
| 12-well culture plates | Fisher | Cat#130185 |
| Sorvall ST 16R bench centrifuge | Thermo Scientific | Cat#75004381 |
| Leica DM6 B upright microscope | Leica | Serial#SN-597425 |
| Leica DMi8 inverted microscope | Leica | Serial#SN-595981 |
| Leica K8 camera | Leica | https://downloads.leica-microsystems.com/K8/Brochure or Flyer/K8_Flyer_EN.pdf |
| Stereo microscope | Nikon | Cat#SMZ1000 |
| Orbital shaker incubator | Scientific Industries | Cat#SI-G100 |

| | | |
|---|---|---|
| Cellometer spectrum image cytometry system | Revvity | https://www.revvity.com/product/cellometer-spectrum-sys1-10x-std-bundle-spectrum-sys1-10x |
| SomnoFlo (Low-flow electronic vaporizer) | Kent Scientific | https://www.kentscientific.com/products/somnoflo/?srsltid=AfmBOoq9FRZZ23Z2WxlWEQKyJHxF0EFnAbW4DKdbb20QqhH_3U8gbpK5 |
| Cryostat CM 1950 | Leica | https://www.leicabiosystems.com/us/histology-equipment/cryostats/leica-cm1950/ |
| Cytek Aurora | Cytek Biosciences | https://cytekbio.com/pages/aurora |
| LS Columns | Miltenyi Biotec | Cat#130-042-401 |
| Hydrophobic barrier pen | Electron Microscopy Sciences | Cat#71312 |
| Superfrost Plus slides | Thermo Fisher Scientific | Cat#12-550-15 |
| Tissue-Tek® OCT compound | Fisher Scientific | Cat#23-730-571 |
| Cellometer slides | Revvity | Cat#CHT4-SD100-014 |
| Insulin syringes | Exel International | Cat#26027 |
| Wound clips | Fisher | Cat#01-804-5 |
| Forane (Isoflurane) | Baxter Healthsciences | Cat#10019-0360-40 |

**Table S2. Base organoid media.**

| **Reagent** | **Final concentration** | **Amount** |
|---|---|---|
| Advanced DMEM/F12 | 1X | 479 mL |
| B27 Supplement | 1X | 10 mL |
| Insulin-transferrin-selenium | 1X | 5 mL |
| Primocin | 100 µg/mL | 1 mL |
| Glutamax | 1X | 5 mL |
| **Total** | | **500 mL** |

**Note:** Can be prepared in advance and stored up to 1 month at 4°C.

**Table S3. Organoid expansion media.**

| Reagent | Stock concentration | Final concentration | Amount |
|---|---|---|---|
| Base organoid media | 1X | 1X | 196.4 mL |
| A83-01 | 500 µM | 500 nM | 200 µL |
| Murine EGF | 50 µg/mL | 50 ng/mL | 200 µL |
| Murine FGF-10 | 100 µg/mL | 100 ng/mL | 200 µL |
| Murine R-spondin1 | 200 µg/mL | 200 ng/mL | 200 µL |
| Murine Noggin | 100 µg/mL | 100 ng/mL | 200 µL |
| Murine Wnt3a | 50 µg/mL | 50 ng/mL | 200 µL |
| Nicotinamide | 0.5 M | 1 mM | 400 µL |
| N2 | 100X | 1X | 2 mL |
| **Total** | | | **200 mL** |

**Note:** Can be prepared in advance and stored up to 2 weeks at 4°C.

#### Videos:

**Video S1.** Representative ultrasound at GD 12 following EEO transplantation into the right uterine horn after epithelial ablation. Left-to-Right scanning shows pregnancy in the EEO-transplanted right uterine horn, with no evidence of pregnancy in the non-transplanted left uterine horn.
